# *In silico* Investigation of Molecular Networks Linking Gastrointestinal Diseases, Malnutrition, and Sarcopenia

**DOI:** 10.1101/2022.03.23.485443

**Authors:** Matti Hoch, Luise Ehlers, Karen Bannert, Christina Stanke, David Brauer, Vanessa Caton, Georg Lamprecht, Olaf Wolkenhauer, Robert Jaster, Markus Wolfien

## Abstract

Malnutrition is a common primary or secondary complication in gastrointestinal diseases. The patient’s nutritional status also influences muscle mass and function, which can be impaired up to the degree of sarcopenia. The molecular interactions in diseases leading to sarcopenia are complex and multifaceted, affecting muscle physiology, the intestine (nutrition), and the liver at different levels. Although extensive knowledge of individual molecular factors is available, their regulatory interplay is not yet fully understood. A comprehensive overall picture of pathological mechanisms and resulting phenotypes is lacking. *In silico* approaches that convert existing knowledge into computationally readable formats can help to unravel such complex systems. We compiled available experimental evidence for molecular interactions involved in the development of sarcopenia into a knowledge base, referred to as the Sarcopenia Map. By including specific diseases, namely liver cirrhosis, and intestinal dysfunction, and considering their effects on nutritional status and blood secretome, we investigated their contribution to the development of sarcopenia. The Sarcopenia Map is publicly available as an open-source, interactive online resource, providing tools that allow users to explore the information on the map and perform *in silico* perturbation experiments.

## Introduction

**Malnutrition** (MN) is a common and characteristic feature of gastrointestinal diseases, such as liver cirrhosis (LC) and intestinal dysfunctions (ID), e.g. short bowel syndrome (SBS), and is connected to high mortality rates ^1^. For LC patients, the prevalence of MN is indicated with up to 90% ^2^; for patients suffering from SBS with around 10 to 40% ^3^. Disease-related MN is closely related to mild, chronic inflammation ^4^. Both MN and inflammation contribute to muscle wasting, which, combined with the loss of muscle function (**sarcopenia**), can negatively influence gastrointestinal diseases. Therefore, sarcopenia is a common secondary phenomenon in LC and ID. The vicious cycle of MN, inflammation, sarcopenia, and the underlying disease itself leads to an unfavorable prognosis for the patient ^5^. The molecular landscape of gastrointestinal disease, nutrition, and muscle metabolism each involves hundreds to thousands of molecules and proteins that form complex molecular communication networks ^6–8^. Linking these processes again increases the complexity many times over. Although the role of many molecules has been elucidated by extensive *in vitro* and *in vivo* experiments, understanding the system as a whole, including all the interactions involved, is a task beyond human capabilities. Therefore, the use of *in silico* approaches, i.e., the conversion of available knowledge into computationally readable formats, can help unravel its complex structure.

**Disease maps** have emerged as web-based knowledge bases summarizing information about molecular interactions in standardized graph representations enabling disease-specific interactive visualizations and *in silico* experiments ^9^. Examples of disease maps include the Atlas of Inflammation Resolution (AIR) ^10^, the Parkinson’s Disease Map ^11^, the Rheumatoid Arthritis Map ^12^, the AsthmaMap ^13^, the Atherosclerosis Map ^14^, or the COVID-19 disease map ^15^. Many of those have been published on **MINERVA**, a web platform that provides interactive visualizations of standardized molecular knowledge graphs ^16^. MINERVA provides automatic annotation, linkage of molecules to external databases, and screening for associated miRNAs, drugs, or chemicals in the graph. In addition, MINERVA allows the development of tools that interact with the map components, making it an excellent framework for web-based disease map presentations. The use of standards such as the systems biology markup language (SBML) ^17,18^ ensures reproducibility, and allows the development of general-purpose tools and automatic annotation, following FAIR principles for scientific data ^19^. By modularizing the interactions into cell type-, tissue-, or process-specific parts, i.e., creating so-called **submaps**, disease maps help to provide an intuitive overview of complex disease mechanisms. These submaps can further be enriched with regulatory interactions from transcription factors, microRNAs (miRNAs) and lncRNAs, or even drugs and chemicals, providing a comprehensive, large-scale **molecular interaction map (MIM)** ^10^. The MIM itself is a directed graph encompassing all interactions that connect the elements, i.e., biological entities in the graph, through activating (positive) or inactivating (negative) causal links. Although the information in disease maps is standardized, the process of creating disease maps, their analysis, and the ultimate search for information is not predetermined and depends mainly on the context, style, and research question.

**Topological analyses** trace causal interactions throughout the network and track the occurrence of nodes ^20,21^. This allows to identify relationships between distant elements, identify hub elements in pathways or provide a weighting of elements regulating a specific phenotype ^22^. Topological methods have also been used to extract core regulatory networks from large-scale networks to investigate mechanisms on a smaller scale ^23^. In addition, topological information has been used to improve the analytical performance of statistical enrichment ^24^ or machine learning approaches ^25^. Topological analysis is less complex but can be problematic in highly interconnected networks. Identifying all paths in larger networks, i.e., all connections between every pair of elements, is a computationally intensive challenge. Consequently, many algorithms focus on identifying only the shortest paths between two nodes in the graph ^26^. Moreover, topological analysis can be highly affected by biases such as (i) misestimating the length of interactions that lack intermediates, (ii) neglecting the biochemical relevance of longer pathways, and (iii) overrepresenting more intensively studied molecules. Nevertheless, they provide a middle ground between ease of implementation and informative power to compare which elements are included in a given pathway and to what extent.

**Boolean models** are much better suited to study actual network mechanisms and to investigate the effects of molecular perturbations. In Boolean models, the **state** of each gene/molecule/phenotype is constrained to be either active (**ON/**1) or inactive (**OFF**/0), defined by specific Boolean rules based on the state of other network elements ^27^. In successive steps, representing a time scale, the state of each element in the map is evaluated based on states during the previous step. Since there is a finite number of possible network states, at some point a **steady state** is reached that is either stable, i.e., remains in one state, or oscillates, i.e., changes infinitely often between several states. The steady state provides useful qualitative information on molecular mechanisms, in particular on circulating regulatory feedback and feedforward loops. Analysis of the number of active states as a function of a given input makes it possible to determine correlations between elements regardless of their distance in the network ^28,29^. This is of great importance in complex processes such as energy metabolism, where the influence of each nutrient must be considered equally at each time point. Moreover, in Boolean models, the computational time increases only proportionally to the complexity of the network, allowing efficient high-throughput analyses. Recently, Montagud et *al*. created personalized Boolean models from clinical data of cancer patients to predict drug targets and validated their results with cell line-specific models ^30^. The development of a Boolean model simulating the influence of nutrition and metabolism on sarcopenia may therefore prove useful in assessing the effects of various physiological and pathological conditions.

We developed an in-depth, standardized, and computationally encoded disease map of the molecular regulatory environment that regulates sarcopenia. We integrated the two disease states, ID and LC, into the map by describing their impact on nutritional and sarcopenic molecular processes. We modularized the map into tissue-specific sub-maps for i) the intestine, i.e., nutrient uptake, its hormonal regulation, and the effects of ID, ii) the liver, i.e., metabolic processes, cytokine secretion, and their alteration in LC, iii) and the muscle, i.e., molecular regulation of catabolic and anabolic muscular processes leading to sarcopenia. In addition, we converted the underlying interaction network into a Boolean model and validated the model by investigating how molecular perturbations are perpetuated in the model. We integrated our methods into an interactive MINERVA tool suite allowing researchers to explore the information on the maps, identify interaction pathways, and perform *in silico* perturbation experiments. With these tools, we demonstrate how the map contributes to understanding the complex molecular processes leading to sarcopenia.

## Methods

### Map creation and curation

We screened the PubMed database for published literature focusing on reviews describing the intestinal uptake of nutrients and their metabolism in the liver, hormonal communication between liver and muscle, and regulation of muscle growth and function. Simultaneously, we sought information on the effects of ID and LC on these processes. We collected the information in three Systems Biology Markup Language (SBML)-standardized **submaps** in CellDesigner ^18,31^. To improve clarity and ease curation efforts. Intracellular molecules were enclosed in compartments reflecting the organ while extracellular molecules were placed outside the compartments, either representing molecules in the bloodstream (e.g., nutrients or cytokines) or systemic conditions, such as acidosis or hyperammonemia. This separation enables the distinction between tissue-specific processes and connects them through the intervening communication processes. Figure 1A provides a schematic overview of the map organization ad the hierarchical flow of information through the submaps.

**Figure 1:**
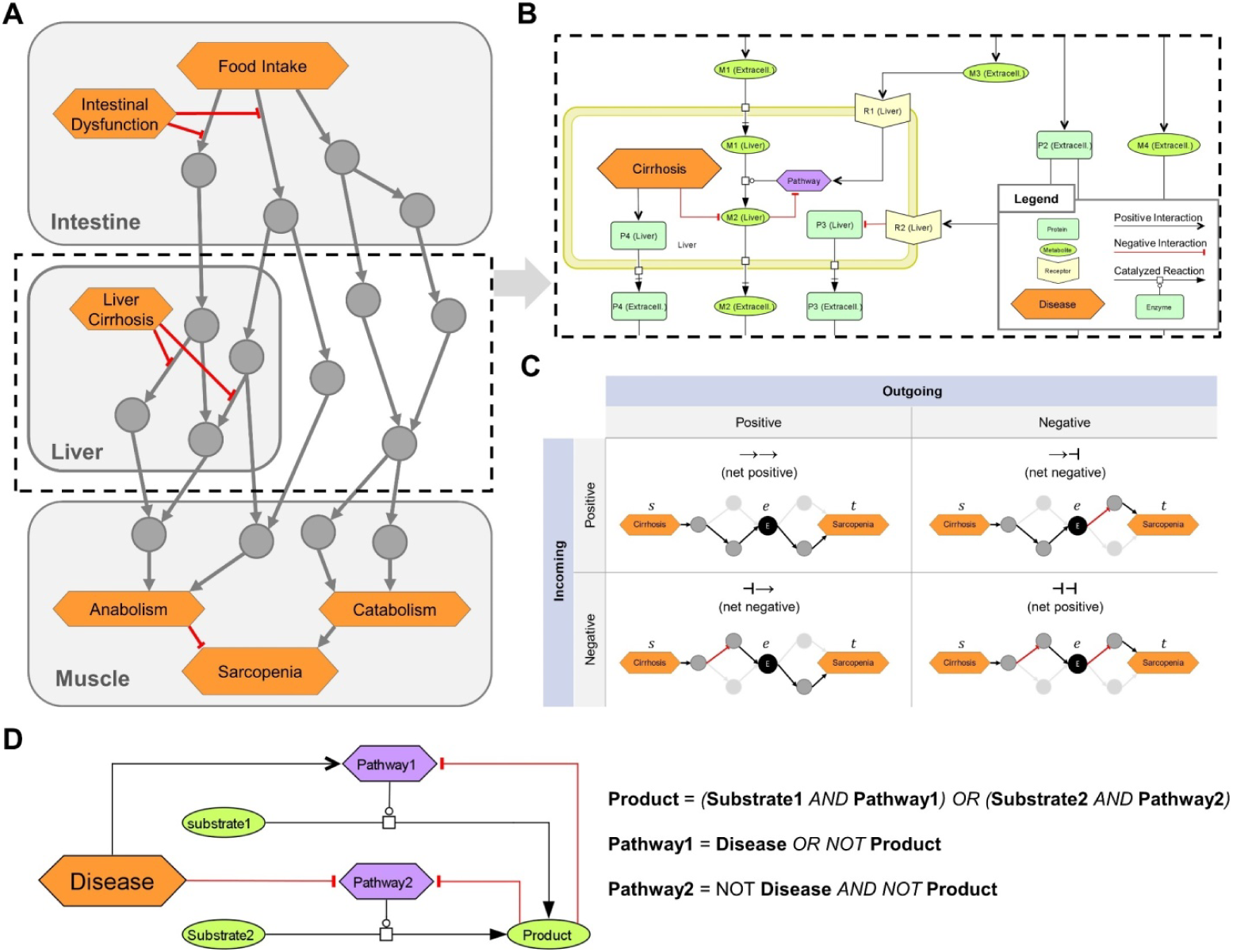
Overview of the logical modeling approaches in the sarcopenia map. (A) Schematic structure of the Sarcopenia Map. (B) SBML schema of tissue-specific compartments (yellow frames) that connected through extracellular elements, e.g., representing hormones or cytokines. (C) Topological analysis of the underlying network. Paths between two elements are analyzed based on their length, type (positive or negative), and passed nodes. Elements are ranked by their inclusions in the identified paths. (D) Creation of Boolean rules that define an element’s state by converting SBML reactions into logical gates.

Different shapes and colors enable intuitive visualizations for various biological or clinical entities, including (i) molecules, such as genes, proteins, or metabolites; (ii) their subclasses, such as receptors and ion channels; (iii) clinical features; (iv) whole pathways. All these entities we collectively refer to as **map elements** (Figure 1B). We connected the elements by SBML-standardized reactions representing their biochemical interaction. We simplified most reactions in the activity flow format, i.e., represented more complex mechanisms (e.g., phosphorylations) as single arrows connecting a source element (e.g., the protein kinase) with a target element (e.g., the protein). This simplification reduces the map content and improves readability while retaining all necessary information. Only enzymatic reactions were retained in the process description format because information about enzymes is necessary for modeling the mechanisms of metabolic regulations. Larger metabolic pathways, e.g., glycolysis, have been combined into a single catalytic reaction leading from the initial reactant (glucose) to the final product (pyruvate), omitting all intermediates. The reaction is catalyzed by a phenotypic element (glycolysis) that represents the metabolic pathway per se. All regulations, e.g., product-feedback-inhibitions or hormonal, were then added as reactions to the phenotype element (exemplarily shown in Figure 1B).

### Directed Interaction Network

We transformed the maps into a single graph (G) consisting of a set of **elements (vertices *V* (*G*))**connected by **interactions (edges *E*(*G*))**. All reactions in the submaps were converted into one or multiple interactions each consisting of two elements that are linked by either upregulation (positive) or downregulation (negative). Therefore, *E*(*G*) is defined as a collection of triples *E* ⊂ (*s* | *r* | *t*)consisting of a source element *s* ∈ *V*, a relation *r* ∈ { −1,1}, and a target element *t* ∈ *V*. All enzymatic reactions in the maps, which catalyze the synthesis of a product *p* from a substrate *s* by an enzyme *e*, were transformed into an edge triplet of (s|1|p), (e|1|p), and (e|−1|s). The latter represents the consumption of the substrate by the enzyme. For reactions with multiple substrates, all substrates were first combined into a complex with edges connecting the substrates and the complex. The complex *c* then acts as the substrate of the reaction triplet. For reactions with multiple products, reaction triplets were generated for each product. A **path** *P*(*G*) of the length *L* ∈ ℕ can be written as the sequence 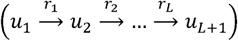 with (*u*_*i*_, *r*_*i*_, *u* _*i* + *1*_) ∈ *E*. The type *T* ∈ {− 1,1} of any *P* is defined as (*r*_1_ · *r*_2_ · … ·*r*_*L*_).

### Topological Model

We identified paths in the MIM using a breadth-first-search (BFS) algorithm, one of the fastest possible solutions in a directed and unweighted graph ^32^. In its standard form, the algorithm enables the search for **shortest paths** (*SP*) between two elements (*u,v*) ∈ *V* as a set of existing paths *P*_*u,v*_ between *u* and *v*, where *L* (*P*_*u,v*_) is minimized. To identify more paths between *u* and *v*, we adapted the BFS algorithm to stop at already visited interactions instead of visited elements. The set of all identified paths or SPs that connect at least two specified elements, we call a **pathway** in the graph. To determine the role of an element *e* in the pathway of *u* and *v*, we filtered the paths between *u* and *v* by those that go through *e*. In addition, the filtered paths were split into two subpaths, *P*_*s,e*_, from *s* to *e* (incoming), and *P*_*e,t*_, from *e* to *t* (outgoing), with *T*(*P*_*s*_,_*t*_) = *T*(*P*_*s,e*_) · *T*(*P*_*e,t*_) and *L*(*P*_*s*_,_*t*_) = *L*(*P*_*s,e*_) + *L*(*P*_*e,t*_) (Figure 1C). This separation of paths provides us with information on i) the ratio of positive and negative paths between *s* and *t*, which *e* is involved in, and ii) the ratio of how *e* is regulated by *s* and how *e* regulates *t*. Repeating this analysis for other elements in the MIM compares their role in the investigated pathway.

### Boolean Model

Based on the interactions in the submaps, we defined a Boolean rule for each element that specifies how its state (either ON or OFF) is defined by the state of other elements (inputs) represented by logical gates (NOT, OR, or AND). A Boolean rule may consist of multiple gates, which may be nested. When a reaction requires multiple elements to be active, such as in enzymatic reactions or the formation of complexes, these elements are represented by AND gates. Any negative input, such as from a disease or negative feedback, is integrated as a NOT gate. In general, all logic gates must be satisfied for an element to be ON, with disease inputs taking precedence. An exception, however, is the glycogen element in the model, whose state we represent as an integer that increases by 1 at each step at which the element’s Boolean rule is satisfied and decreases by 1 when it is not. As long as its state is greater than zero, it is treated as an ON input to other elements. In this way, we can simulate the construction of a storage and its subsequent use, even after its inputs subsided. Finally, we defined an initial state of the model in which some elements with no inputs, such as digestive enzymes or transporters, are set to ON. Supplementary File 1 shows the list of map elements with their initial state.

### Sensitivity Analyses

We identified the correlations between two elements based on the dependency of their activity. The principle behind this methodology has been described by Helikar et *al*. in 2009 ^29^. The activity of an element is defined as the percentage *p* of active states of the model during a range of *n* observed steps (*n* =100 by default). We perturbed a source element *s* either through a set activity or through inhibition. Setting the activity of an element means changing its state to OFF and then to ON at every k-th step depending on the activity frequency *p*_*a*_ (*s*) with 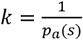. For example, the input activity frequency *p*_*a*_ (*s*) =0.25 refers to a state sequence for *s* of [1-0-0-0-1-0-0-0-1-…]. When perturbing *s* through inhibition, its state is set to OFF for every *k*-th step with 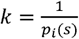 while in all other steps the element behaves normally. Then after performing *n* steps, we measured the activity of the target element *t* as the percentage of steps with ON state. If *t* is the ‘sarcopenia’ phenotype in the muscle, its activity is defined as

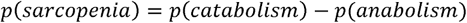

The phenotypes ‘anabolism’ and ‘catabolism’ themselves are defined as:

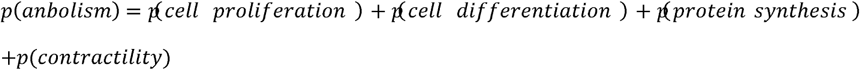

and

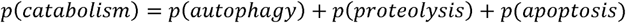

The perturbation experiment was then repeated for different activities of *s*. Finally, the correlation between both elements was analyzed using the ‘pearsonr’ function from the stats module of the scipy python package generating the Pearson correlation coefficient and a two-sided *p*-value.

## Results

We present the Sarcopenia Map as a publicly available, comprehensive knowledge base of experimental evidence for molecular interactions underlying sarcopenia, linking it to ID and LC (Figure 2). In the following, we present (i) the Sarcopenia Map as a knowledge base (ii) its tools to perform *in silico* simulations, and (iii) applications and validations of the underlying computational model. We provide examples of how the tools help researchers analyzing disease mechanisms by investigating the molecular interactions connecting nutrition, gastrointestinal diseases, and sarcopenia.

**Figure 2:**
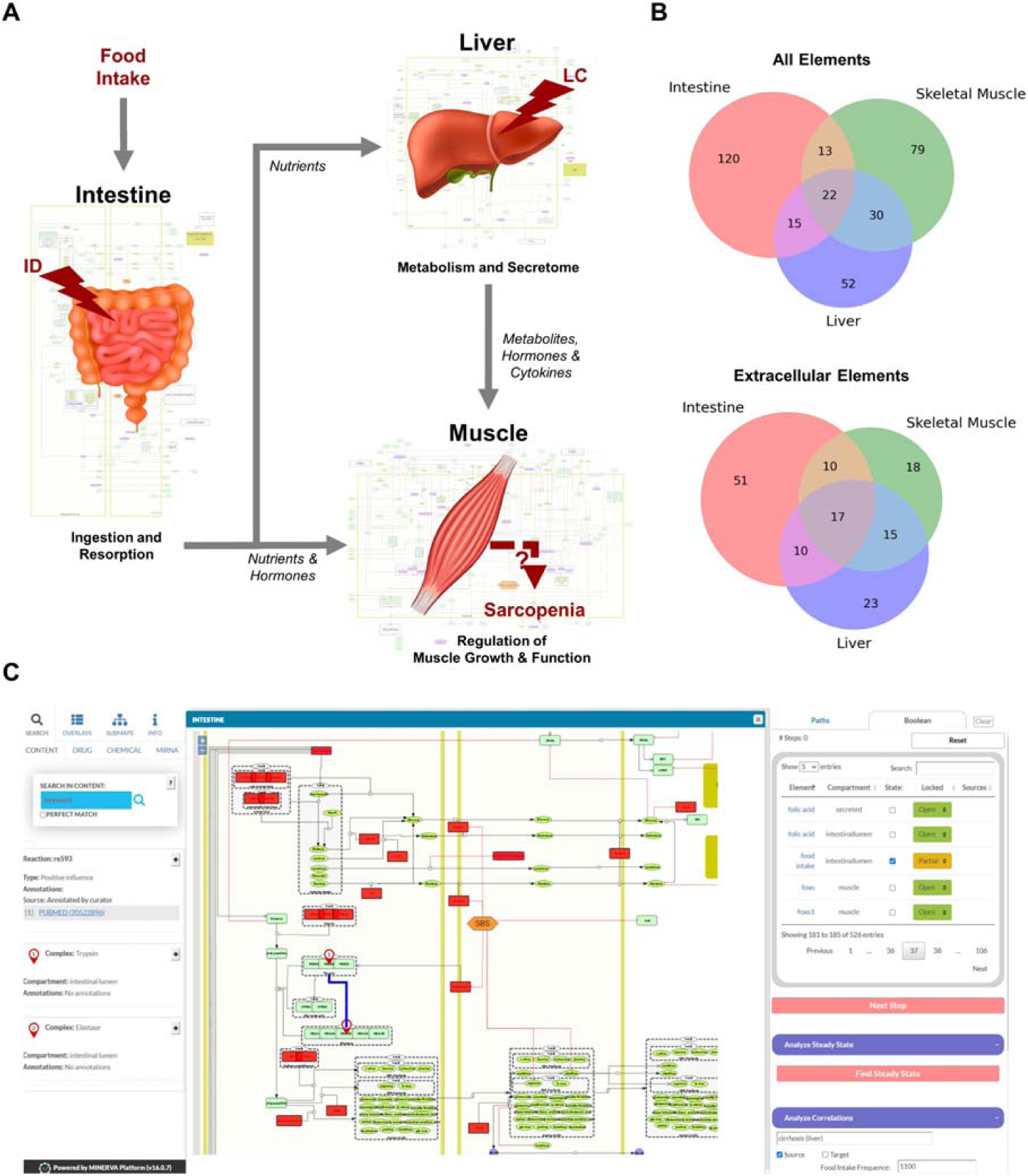
Overview of the hierarchical organization of the Sarcopenia map. (A) We summarized information on molecular interactions related to sarcopenia from the literature into three tissue-specific submaps. In addition, we integrated the effects of liver cirrhosis (LC) and intestinal dysfunction (ID) on these molecular processes. (B) Venn Diagrams of element distributions between the different tissues. (C) The interactive user interface of the Sarcopenia Map enables the exploration of information as well as simulations of molecular perturbations.

### A Knowledge Base of Regulatory Mechanisms in Sarcopenia

We have compiled findings from the scientific literature into three standardized, tissue-specific submaps (Figure 2A). The submaps summarize the processes in each tissue as SBML-standardized molecular networks (Supplementary Figure 1, Supplementary Figure 2, and Supplementary Figure 3). Figure 2B shows the distribution of elements between submaps. Their overlap is more dominant for extracellular elements, which is to be expected since they represent secreted molecules that communicate between compartments. The highest number of tissue unique elements is found in the submap of the intestine, as it contains food components as well as digestive enzymes and transporters.

MINERVA provides features to explore elements in the map and targets of specific drugs, miRNAs, or chemicals. The submaps are publicly accessible and can be downloaded in various formats (e.g., SBML, .svg, or .pdf). All elements and interactions in the maps are annotated with references to public databases or scientific literature (e.g., PubMed). The Sarcopenia Map comes with an interactive tool that we developed to allow users to explore interaction paths in the sarcopenia map through topological analysis and to perform *in silico* simulations with the Boolean model in an easy-to-use interface (Figure 2C).

### A Tool for Topological and Boolean Analyses

The first part of the map tool provides network topology analyses to investigate, which elements are included in filtered interaction pathways between specific elements to explore their underlying molecular regulatory mechanisms. It provides an overview of the elements included in filtered interaction paths between specific elements to explore their underlying molecular regulatory mechanisms. Users select a source element (“*From*”) and a target element (“*To*”) whose interaction paths are to be identified in the MIM (Figure 3A). In addition, another element can be specified to show only the interactions that pass “*Through*” that element. The output will be a table that shows all identified paths, their length, the total impact on the target, and the individual steps within the path, including their interactions (Figure 3B). In addition, a bar chart lists the percentage of these paths, in which each element occurs, separated into positive and negative paths (Figure 3C). However, due to the limitations of topological models, assumptions about functional relationships should not be inferred from the distribution of positive and negative interaction paths alone. Nevertheless, they provide an intuitive overview of the design of molecular pathways and the flow of information.

**Figure 3:**
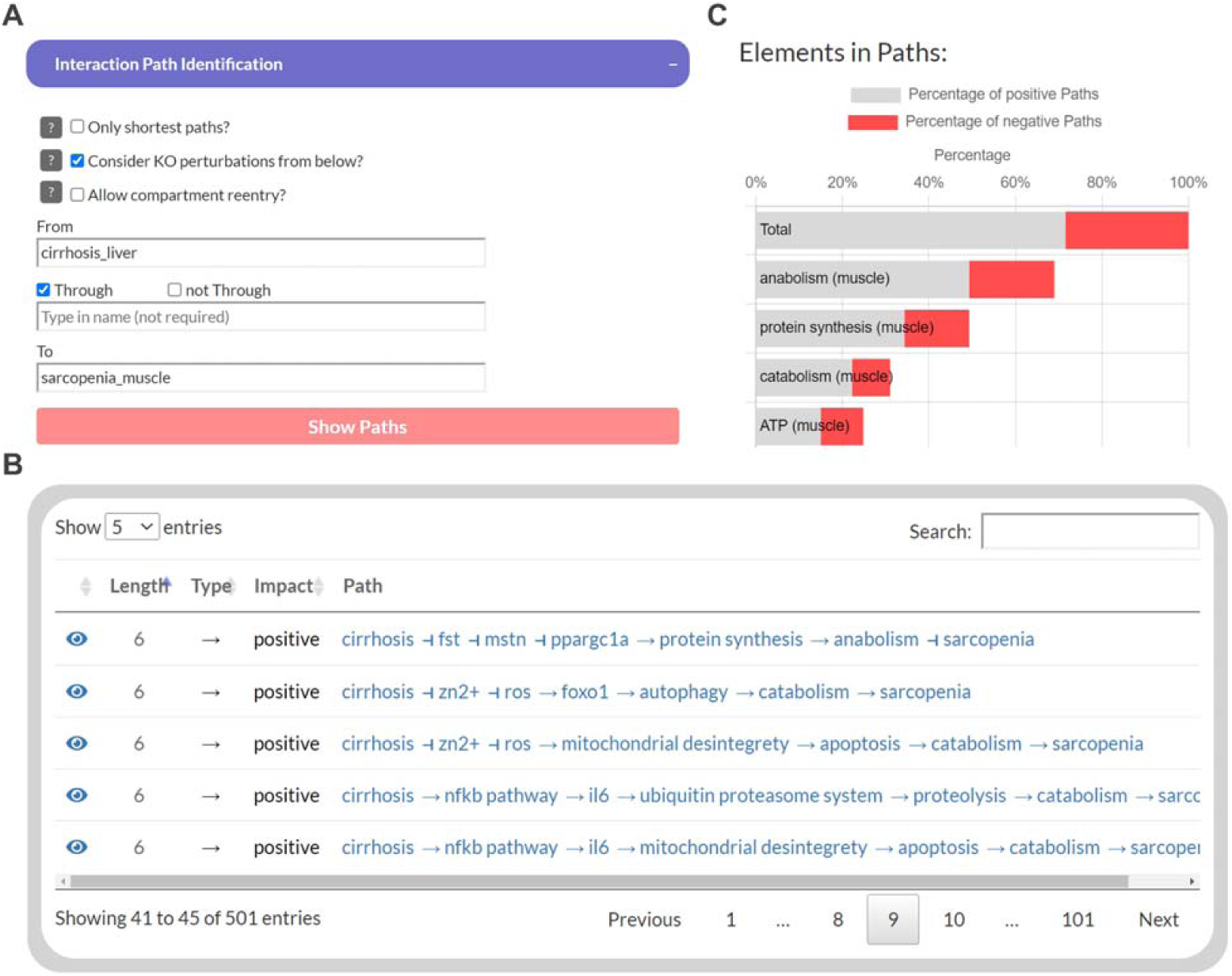
The user interface to identify interaction paths between selected elements in the Sarcopenia map. For selected elements (A), their interaction pathways are listed in a table (B). Additionally, elements along the pathways are ranked by their percentages of appearance separated by the type of interaction (C).

In addition to topological analysis, we created a Boolean model of the sarcopenia map and integrated functions into the map tool to observe individual steps as colored overlays on the maps, identify stable states, and analyze correlations between elements. Supplementary Figure 4 illustrates the activity of elements during the oscillating steady state, which evolves from the default input state of the map and simulates permanently active food input. It can be seen that most of the elements that change state during steady state are metabolites and metabolic enzymes. Feedback loops of metabolic pathways are responsible for this behavior, as shown by the example of glucose metabolism in the liver. As food intake is constantly ON, so is extracellular glucose, leading to a constant oscillation between glycolysis and glycogen synthesis. Feedback loops are essential for reversible reactions in Boolean models because without them, the ON signal would be constantly active in both elements, even if the external signal was OFF. Once we set the food intake to OFF and iterate forward, the new resulting steady state is stable, i.e., it does not fluctuate and has no active metabolites (not shown).

The tool also enables the identification of correlations between elements in the map based on a user-defined state of the model. For a selected source element, its activity is iteratively changed from 0 to 1 in 0.05 increments, and for each, the activity of all other elements is measured for 100 steps. A resulting table presents the Pearson correlation coefficient for all elements and also shows the activity correlation plot. However, in Boolean models, it is difficult to keep track and rank the elements that have transmitted signals throughout the network, since their execution involves hundreds of steps with multiple feedback loops throughout the network. To find a feasible solution for this problem, our tool ranks all elements in the network according to their correlation with the source and destination elements. Elements that correlate with both simultaneously are most likely responsible for transmitting the signals. In addition, based on the type of correlation (positive or negative), we can investigate the role of the transmitting element, i.e., whether inhibition/activation of an inhibitory/activating signal has occurred or *vice versa*.

### Validating and Testing the Boolean Model

To validate the Boolean model, we studied the behavior of the carbohydrate system under different nutrient conditions, i.e., activities of the food intake element. Since carbohydrate metabolism is a tightly regulated system and the central part of the energy cycle that controls muscle function, we need to ensure that in our model carbohydrate levels respond to perturbations similarly to physiological conditions. We tested the model by measuring the response of glucose and glycogen to altered nutritional stimuli. Figure 4A shows the extent of hepatic glycogen storage (blue dots) in response to an increasingly number of steps with an active food intake (y-axis, black dots). As expected, an increased value of hepatic glycogen and an increasingly prolonged decrease after reaching the maximum value were observed when food intake is switched off. In addition, blood glucose (red dots) is continuously active as long as food intake occurs and is oscillating during glycogen depletion. These results show that our model can simulate the conversion of glycogen to glucose and its release into the bloodstream in fasting situations.

**Figure 4:**
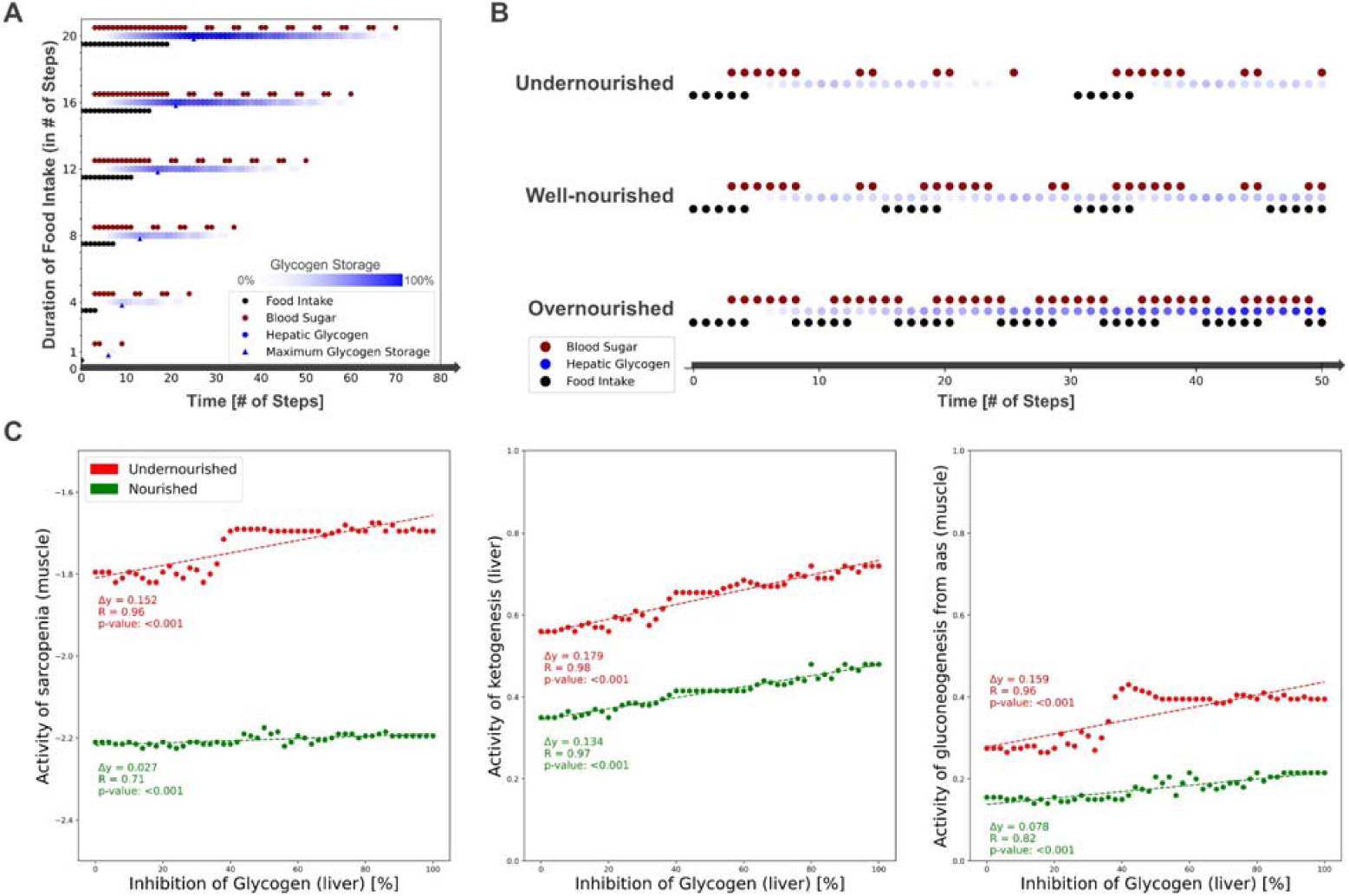
Simulating the carbohydrate availability under different nutritional inputs and investigating the impact of glycogen deficiency. (A) Level of hepatic glycogen and active states of extracellular glucose dependent on the duration of food intake. (B) Level of hepatic glycogen and active states of extracellular glucose in three defined combinations of ON and OFF ‘food intake’ as simulations of different nutritional states. (C) Impact of glucose deficiency on sarcopenia, ketogenesis in the liver, and glucogenesis from amino acids in the muscle depending on the nutritional state.

Next, we measured carbohydrate behavior again, but with different combinations of ON and OFF food intake, representing changing frequency and quantity, but not quality, of diet. From these, we identified three specific **nutrition states** which will act as input for the model to simulate (patho-)physiological behavior. In Figure 4B we show the carbohydrate behavior of the selected nutrition states: (i) undernourished, i.e., long fasting periods with full depletion of glycogen storage between food intake, (ii) well-nourished, with continuous glycogen storage, and (iii) overnourished, with continuously increasing glycogen. The carbohydrate metabolism is a key pathway in connecting gastrointestinal disease with sarcopenia. In clinical settings, glucose supplementation has been shown to reduce muscle mass loss, while glycogen depletion in LC patients has been identified as a major cause of the development of sarcopenia ^33,34^. Therefore, we investigated how a deficiency in glycogen reserves impacts glucose substitution and muscle phenotypes under the selected nutrition states (Figure 4C). Most noticeably, in our model, glycogen inhibition exhibit effects only in well-nourished and undernourished states. At overnourishment, no correlation, and during well-nourishment only a low correlation is visible because extracellular glucose is continuously substituted and, thus, does not necessarily rely on refill from the hepatic glycogen storages. We see that glycogen deficiency positively correlates with sarcopenia, with higher base values and higher regression coefficients in the undernourished than in the well-nourished state. A similar correlation is visible for the ‘ketogenesis’ phenotype in the liver and the ‘glucogenesis from amino acids’ in the muscle, both physiological responses to hypoglycemic states ^34,35^.

Next, we investigated the correlations between activities of LC and ID on the muscle phenotypes ‘anabolism’, ‘catabolism’, and ‘sarcopenia’ dependent on the nutritional state (Figure 5). In both diseases, we see a strong positive correlation with catabolism (blue) and a negative correlation with anabolism (red). Thus, both disease states also correlate positively with sarcopenia. No significant differences are seen between the nutritional states. However, we see a lower contribution of both diseases to anabolism in the undernourished state compared to the other states. Presumably, this is due to the generally lower activity of anabolism in the undernourished state. In general, the correlation tends to be constant in the overnourished state, whereas the correlations diverge more in the nourished and undernourished states. In these two groups, we see a strong increase in catabolism and sarcopenia activity at low LC activities. This may suggest that in malnourished states, even a low LC disease severity has a large effect on muscle phenotypes. In ID, phenotypes show an almost plateau-like behavior at low disease activities (<0.5) and only then start to increase. This is to be expected because at low frequencies of food intake, the effects of the lower activities of ID, which are mainly related to food resorption, are minimal.

**Figure 5:**
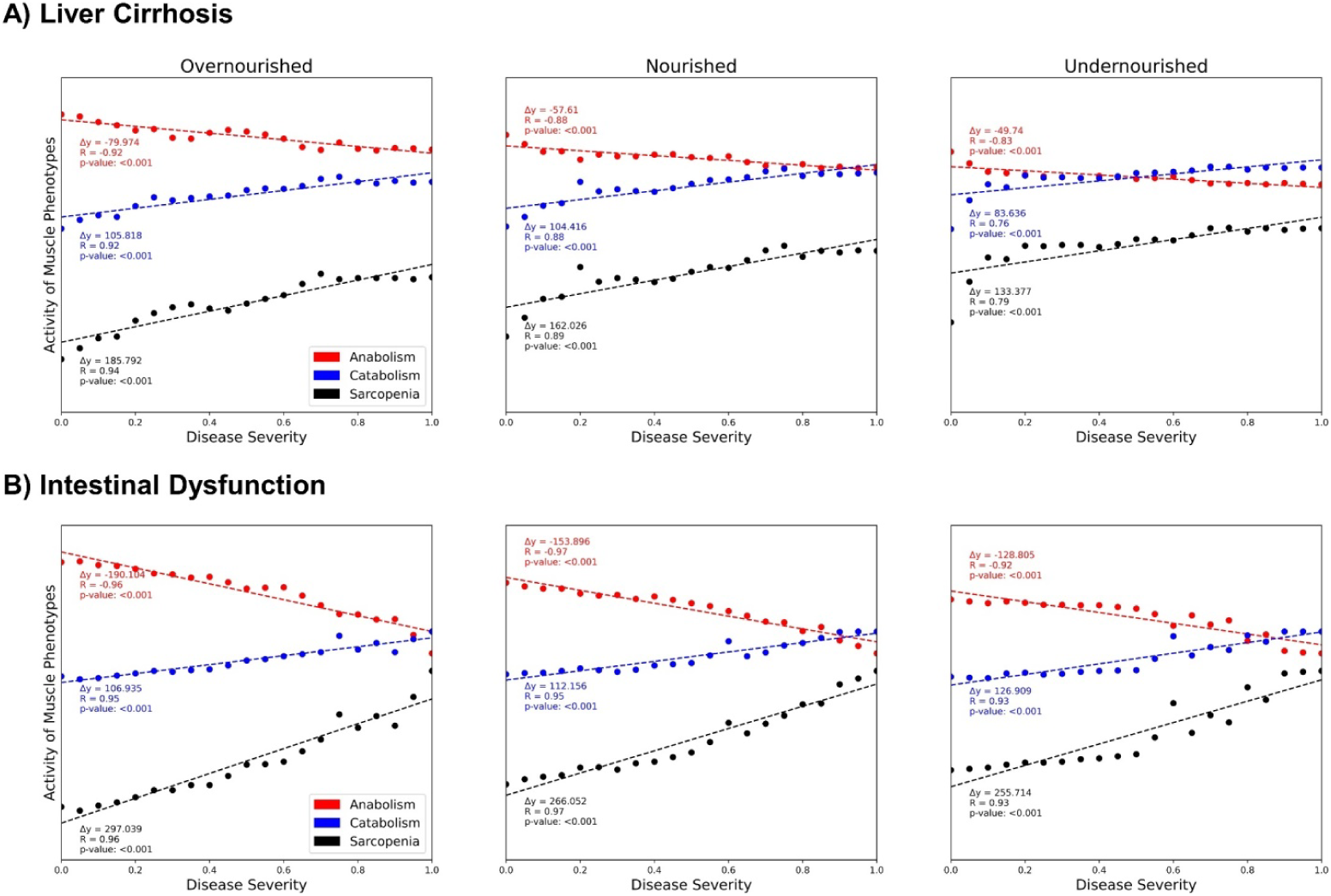
Correlation between the activity of muscle phenotype elements in response to defined activities of liver cirrhosis (A) and intestinal dysfunction (B) in three different nutritional states. Each point represents an experiment in which, starting from an initial state, the state of each element in the network was recalculated 100 times (= steps). The activity of an element is defined as the frequency of its active state among all steps. The activity of anabolism and catabolism is the cumulative activity of all corresponding phenotypes, while the activity of sarcopenia is the difference between catabolism and anabolism.

## Discussion

As scientific knowledge increases so does awareness of the complexity of the molecular mechanisms that regulate biological processes. Gastrointestinal diseases are regulated through complex, interconnected networks in multiple cell types, tissues, and organs ^36^. Muscle growth and function are tightly regulated processes to keep the body functioning in different dietary situations ^5,8^. Therefore, various nutrients and hormones are involved in regulating muscle activity, which complicates the search for the causes of dysregulations, such as sarcopenia. Moreover, the function of individual molecules in these processes can change depending on environmental conditions such as the activity of other elements. Experimental setups are usually unable to simultaneously mimic the complex interactions between different metabolic pathways in various cells, compartments, or tissues. The complex of molecular network linking gastrointestinal diseases, malnutrition and sarcopenia motivate the use of *in silico* methods.

We established the Sarcopenia Map to bring the complex molecular interaction pathways in sarcopenia into a comprehensive, standardized and reproducible format. The Sarcopenia Map is a knowledge base that (i) gathers molecular information, including links to external databases, (ii) is a platform to analyze experimental data on the map, and (iii) allows *in silico* experiments.

By analyzing the topology of a graph representing the large-scale molecular network, users can identify interaction paths for further modeling and simulation. The Boolean model enables the simulation of molecular networks and observe how different activities of molecules could reflect changes in the system as a whole. We showed that our model can be used to simulate molecular mechanisms by successfully reproducing existing knowledge.

We provide the community with a free-to-use platform to support nutrition research in developing or validating new hypotheses. While our work focused on the effects of gastrointestinal diseases such as LC or ID on sarcopenia, the map itself provides a comprehensive knowledge base linking nutrition and muscle metabolism that can be useful for other research areas. The hierarchical format of the map and the standardized representation of molecular interactions facilitate extension to other related diseases or integration of new information, such as malnutrition in relation to other tissues, in the future.

Disease maps are community resources, and MINERVA provides tools for the community to expand these maps collaboratively. We encourage researchers using the sarcopenia map to support open science by sharing scientific results to improve the content of the map.

## Author contributions

OW, RJ, GL, and MW conceptualized and supervised the project. MH, LE, KB, and MW supervised parts that included literature research, curation of content, and layout of submaps. CS, DB, and VC performed literature research and designed the submaps. MH created the model, developed the tools, and performed the analyses. LE supervised both, model design and interpretation of results, in the medical context. MH and LE prepared the initial version of the manuscript. OW, RJ, GL, KB and MW critically evaluated the manuscript and the results. All authors contributed to the scientific content and helped writing the text. All authors approved of the submitted version.

## Data and materials availability

The Sarcopenia Map is available as an open-source resource at https://www.sbi.uni-rostock.de/minerva/index.xhtml?id=SarcopeniaMap

## Supplementary Figures

**Supplementary Figure 1:**
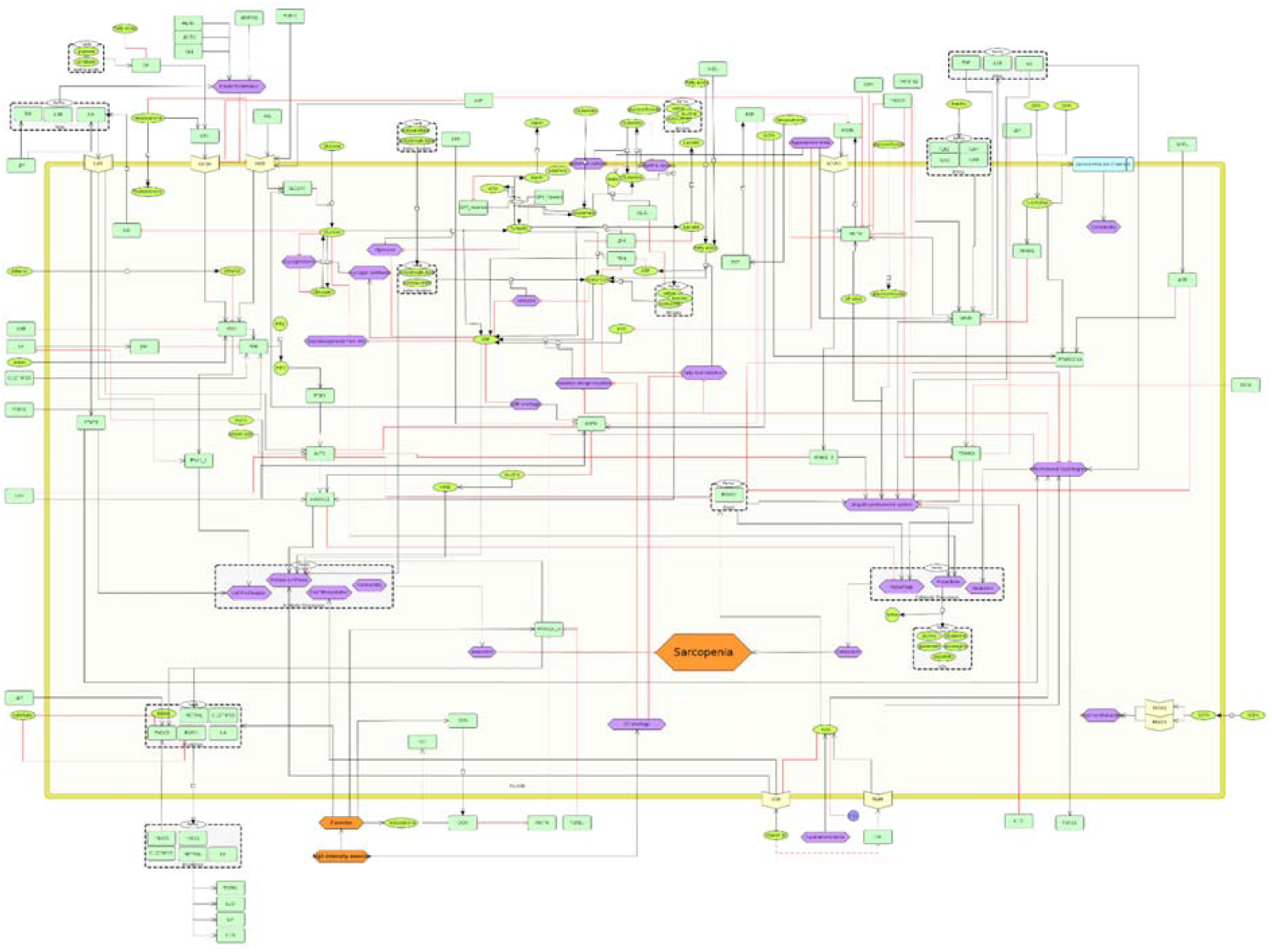
SBML-standardized submap of muscle-specific processes involved in the regulation of sarcopenia. The map includes signaling pathways of hormones, cytokines, and metabolites on muscle anabolism (left) and catabolism (right), thus regulating the development of sarcopenia (orange).

**Supplementary Figure 2:**
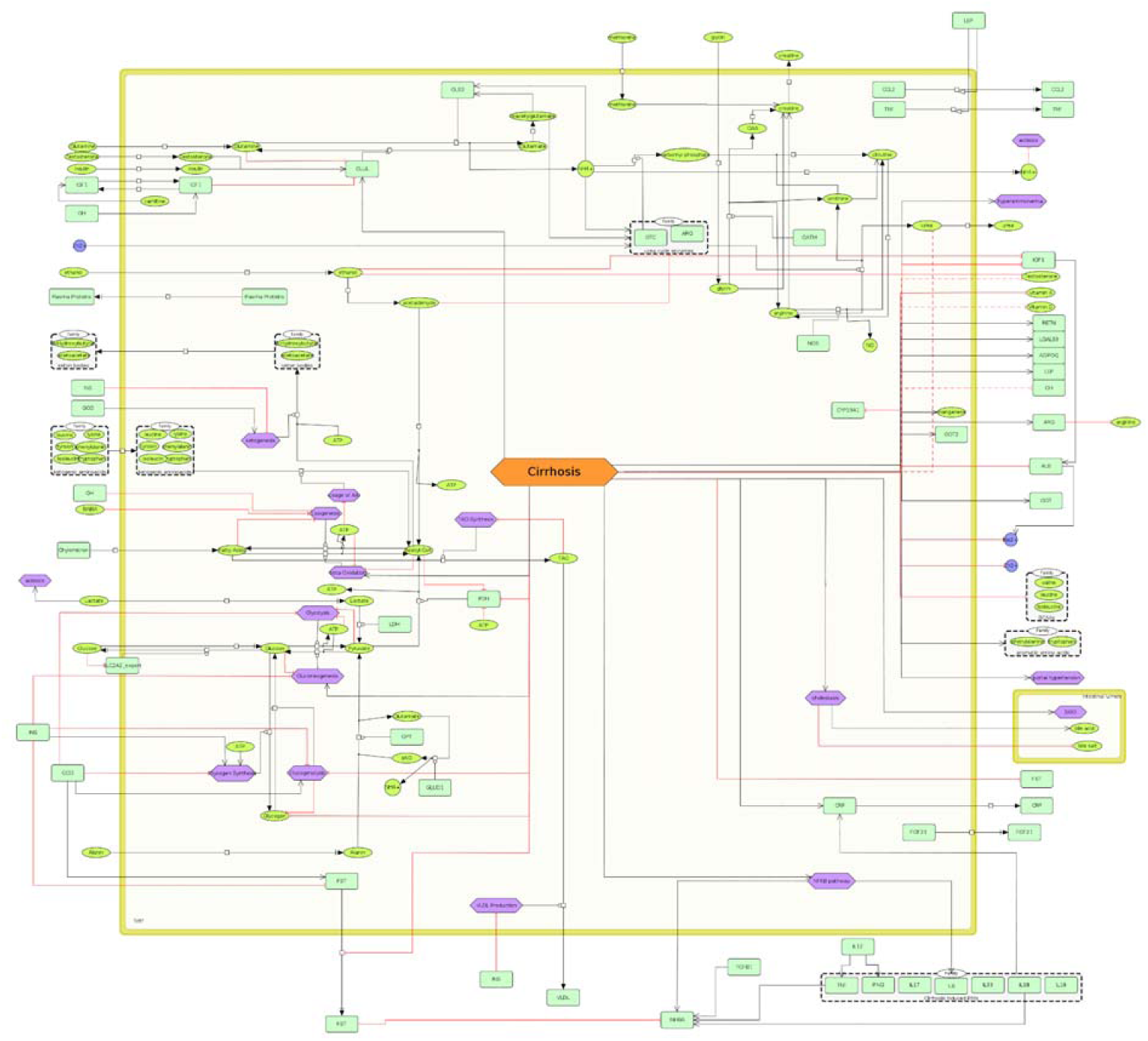
SBML-standardized submap of liver-specific processes involved in the regulation of sarcopenia. The map contains information on metabolic processes (left), secreted hormones or metabolites (right), and their alterations in liver cirrhosis (orange).

**Supplementary Figure 3:**
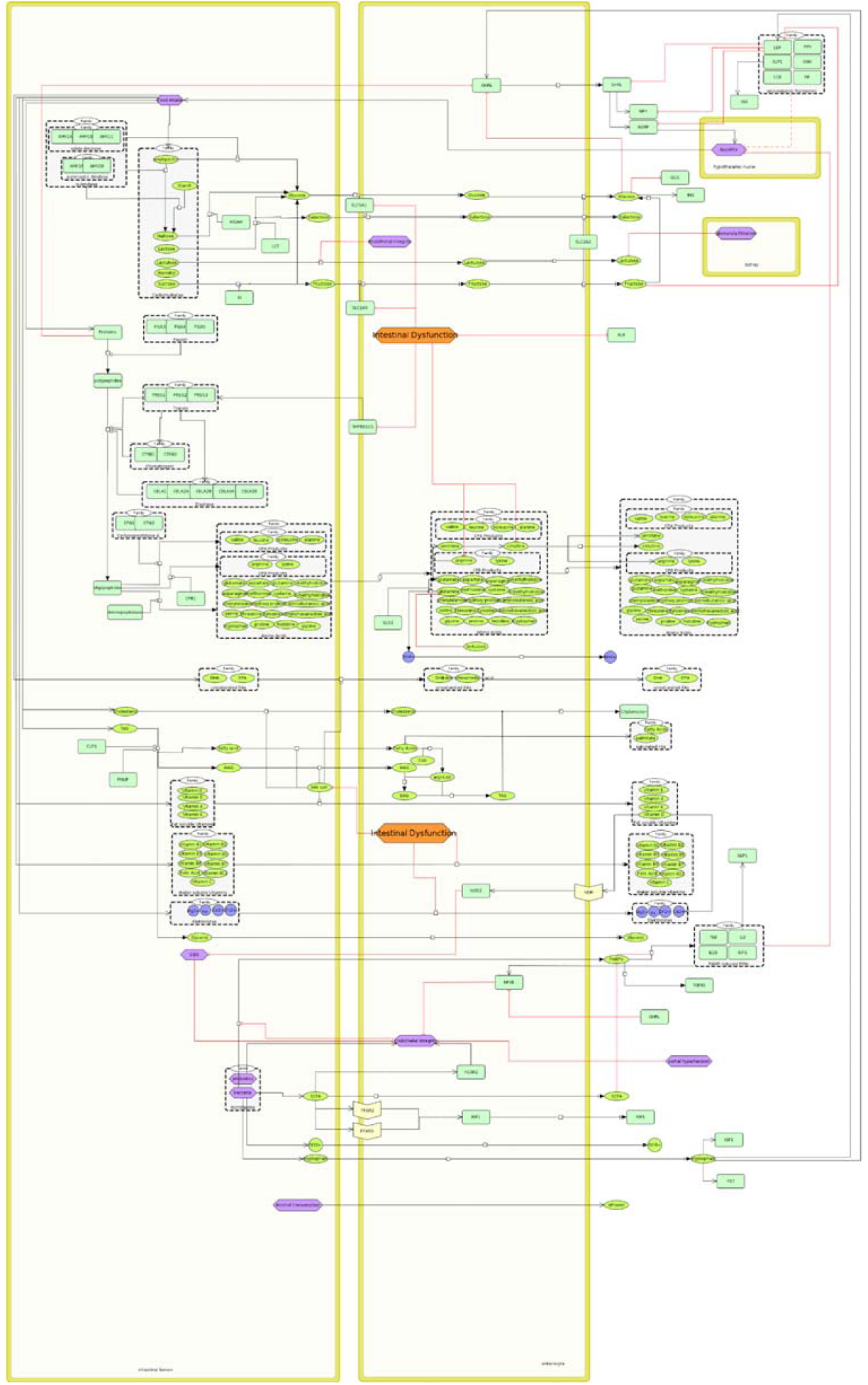
SBML-standardized submap of Gut-specific processes involved in the regulation of sarcopenia. The map contains information on nutrient resorption, secretion of hormones, and their alterations in short bowel syndrome (SBS, orange).

**Supplementary Figure 4:**
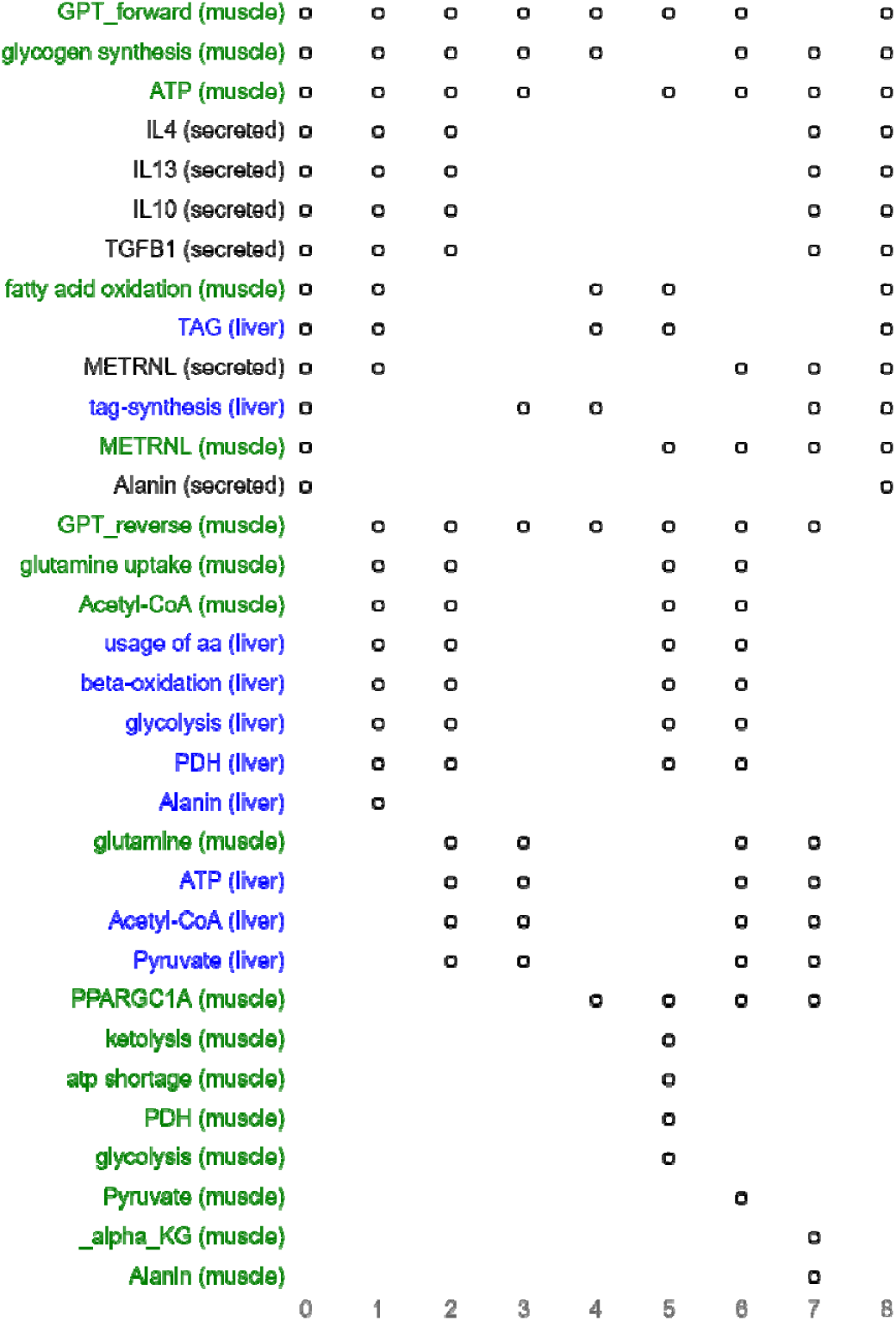
Steady state of the Boolean model with a constantly active ‘food intake’. Each dot represents an active element (y-axis) in the respective step (x-axis) during the steady state. In the last step the original state is reached, and thus the sequence iterates infinitely.

## References

1. M, N. et al. Systematic review with meta-analysis: Nutritional screening and assessment tools in cirrhosis. Liver Int. 40, 664–673 (2020).

2. X, T. et al. A Systematic Review of Medical Nutrition Therapy Guidelines for Liver Cirrhosis: Do We Agree? Nutr. Clin. Pract. 35, 98–107 (2020).

3. Siddiqui, M. T., Al-Yaman, W., Singh, A. & Kirby, D. F. Short-Bowel Syndrome: Epidemiology, Hospitalization Trends, In-Hospital Mortality, and Healthcare Utilization. J. Parenter. Enter. Nutr. 45, 1441–1455 (2021).

4. F, M. & L, V. Disease-Related Malnutrition and Sarcopenia as Determinants of Clinical Outcome. Visc. Med. 35, 282–290 (2019).

5. L, E. et al. Preclinical insights into the gut-skeletal muscle axis in chronic gastrointestinal diseases. J. Cell. Mol. Med. 24, 8304–8314 (2020).

6. Barabási, A. L., Menichetti, G. & Loscalzo, J. The unmapped chemical complexity of our diet. Nat. Food 2019 11 1, 33–37 (2019).

7. Tripathi, A. et al. The gut–liver axis and the intersection with the microbiome. Nat. Rev. Gastroenterol. Hepatol. 2018 157 15, 397–411 (2018).

8. Egan, B. & Zierath, J. R. Exercise Metabolism and the Molecular Regulation of Skeletal Muscle Adaptation. Cell Metab. 17, 162–184 (2013).

9. Mazein, A. et al. Systems medicine disease maps: community-driven comprehensive representation of disease mechanisms. npj Syst. Biol. Appl. 4, 21 (2018).

10. Serhan, C. N. et al. The Atlas of Inflammation Resolution (AIR). Mol. Aspects Med. 74, 100894 (2020).

11. Fujita, K. A. et al. Integrating Pathways of Parkinson’s Disease in a Molecular Interaction Map. Mol. Neurobiol. 49, 88–102 (2014).

12. Singh, V. et al. Computational Systems Biology Approach for the Study of Rheumatoid Arthritis: From a Molecular Map to a Dynamical Model. Genomics Comput. Biol. 4, 100050 (2018).

13. Mazein, A. et al. AsthmaMap: An expert-driven computational representation of disease mechanisms. Clin. Exp. Allergy 48, 916–918 (2018).

14. Parton, A., McGilligan, V., Chemaly, M., O’Kane, M. & Watterson, S. New models of atherosclerosis and multi-drug therapeutic interventions. Bioinformatics 35, 2449–2457 (2019).

15. Ostaszewski, M. et al. COVID19 Disease Map, a computational knowledge repository of virus–host interaction mechanisms. Mol. Syst. Biol. 17, e10387 (2021).

16. Gawron, P. et al. MINERVA—a platform for visualization and curation of molecular interaction networks. npj Syst. Biol. Appl. 2, 16020 (2016).

17. Hucka, M. et al. The systems biology markup language (SBML): A medium for representation and exchange of biochemical network models. Bioinformatics 19, 524–531 (2003).

18. Keating, S. M. et al. <scp>SBML</scp> Level 3: an extensible format for the exchange and reuse of biological models. Mol. Syst. Biol. 16, e9110 (2020).

19. Wilkinson, M. D. et al. The FAIR Guiding Principles for scientific data management and stewardship. Sci. Data 2016 31 3, 1–9 (2016).

20. Koutrouli, M., Karatzas, E., Paez-Espino, D. & Pavlopoulos, G. A. A Guide to Conquer the Biological Network Era Using Graph Theory. Front. Bioeng. Biotechnol. 8, 34 (2020).

21. Janjić, V. & Pržulj, N. Biological function through network topology: a survey of the human diseasome. Brief. Funct. Genomics 11, 522–532 (2012).

22. Hoch, M. et al. Network-and Enrichment-based Inference of Phenotypes and Targets from large-scale Disease Maps (Accepted for publication in npj Systems Biology and Applications). bioRxiv (2022) doi:10.1101/2021.09.13.460023.

23. Khan, F. M. et al. Unraveling a tumor type-specific regulatory core underlying E2F1-mediated epithelial-mesenchymal transition to predict receptor protein signatures. Nat. Commun. 8, (2017).

24. Zito, A. et al. Gene Set Enrichment Analysis of Interaction Networks Weighted by Node Centrality. Front. Genet. 12, 577623 (2021).

25. Liu, H. et al. Predicting effective drug combinations using gradient tree boosting based on features extracted from drug-protein heterogeneous network. BMC Bioinformatics 20, 645 (2019).

26. Klamt, S. & von Kamp, A. Computing paths and cycles in biological interaction graphs. BMC Bioinformatics 10, 181 (2009).

27. Saadatpour, A. & Albert, R. Boolean modeling of biological regulatory networks: A methodology tutorial. Methods 62, 3–12 (2013).

28. Helikar, T. et al. The Cell Collective: Toward an open and collaborative approach to systems biology. BMC Syst. Biol. 6, 1–14 (2012).

29. Helikar, T. & Rogers, J. A. ChemChains: a platform for simulation and analysis of biochemical networks aimed to laboratory scientists. BMC Syst. Biol. 3, 58 (2009).

30. Montagud, A. et al. Patient-specific Boolean models of signalling networks guide personalised treatments. Elife 11, (2022).

31. Funahashi, A., Morohashi, M., Kitano, H. & Tanimura, N. CellDesigner: a process diagram editor for gene-regulatory and biochemical networks. BIOSILICO 1, 159–162 (2003).

32. Liu, Y., Wei, X., Chen, W., Hu, L. & He, Z. A graph-traversal approach to identify influential nodes in a network. Patterns (New York, N.Y.) 2, 100321 (2021).

33. Dhaliwal, A. & Armstrong, M. J. Sarcopenia in cirrhosis: A practical overview. Clin. Med. (Northfield. Il). 20, 489–492 (2020).

34. Ebadi, M., Bhanji, R. A., Mazurak, V. C. & Montano-Loza, A. J. Sarcopenia in cirrhosis: from pathogenesis to interventions. J. Gastroenterol. 54, 845–859 (2019).

35. Laffel, L. Ketone bodies: a review of physiology, pathophysiology and application of monitoring to diabetes. Diabetes. Metab. Res. Rev. 15, 412–426 (1999).

36. Holtmann, G., Shah, A. & Morrison, M. Pathophysiology of Functional Gastrointestinal Disorders: A Holistic Overview. Dig. Dis. 35, 5–13 (2017).

