## Supplementary figures and images for "*In silico* Investigation of Molecular Networks Linking Gastrointestinal Diseases, Malnutrition, and Sarcopenia"

### Supplementary Figure 1

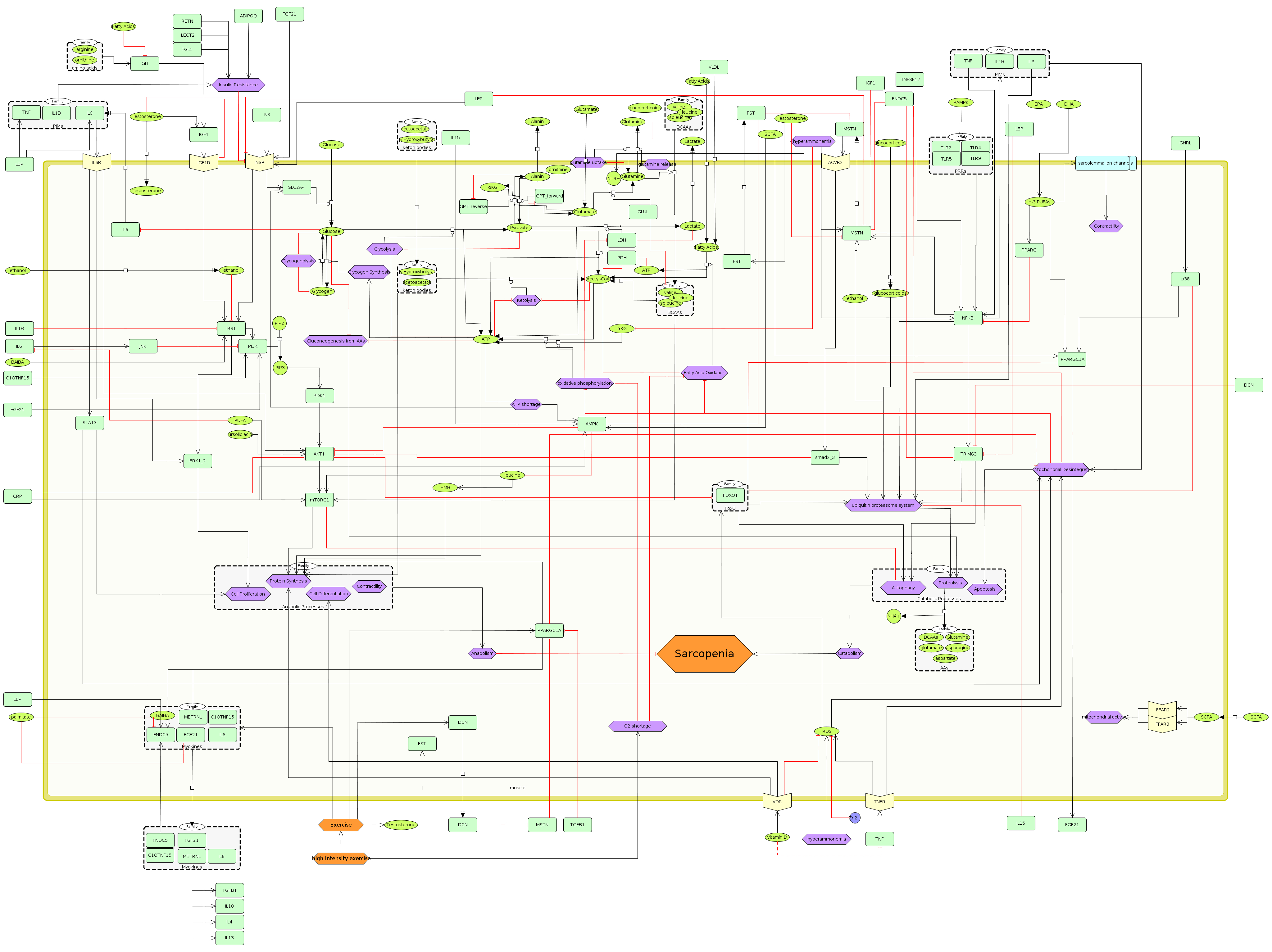

### Supplementary Figure 2

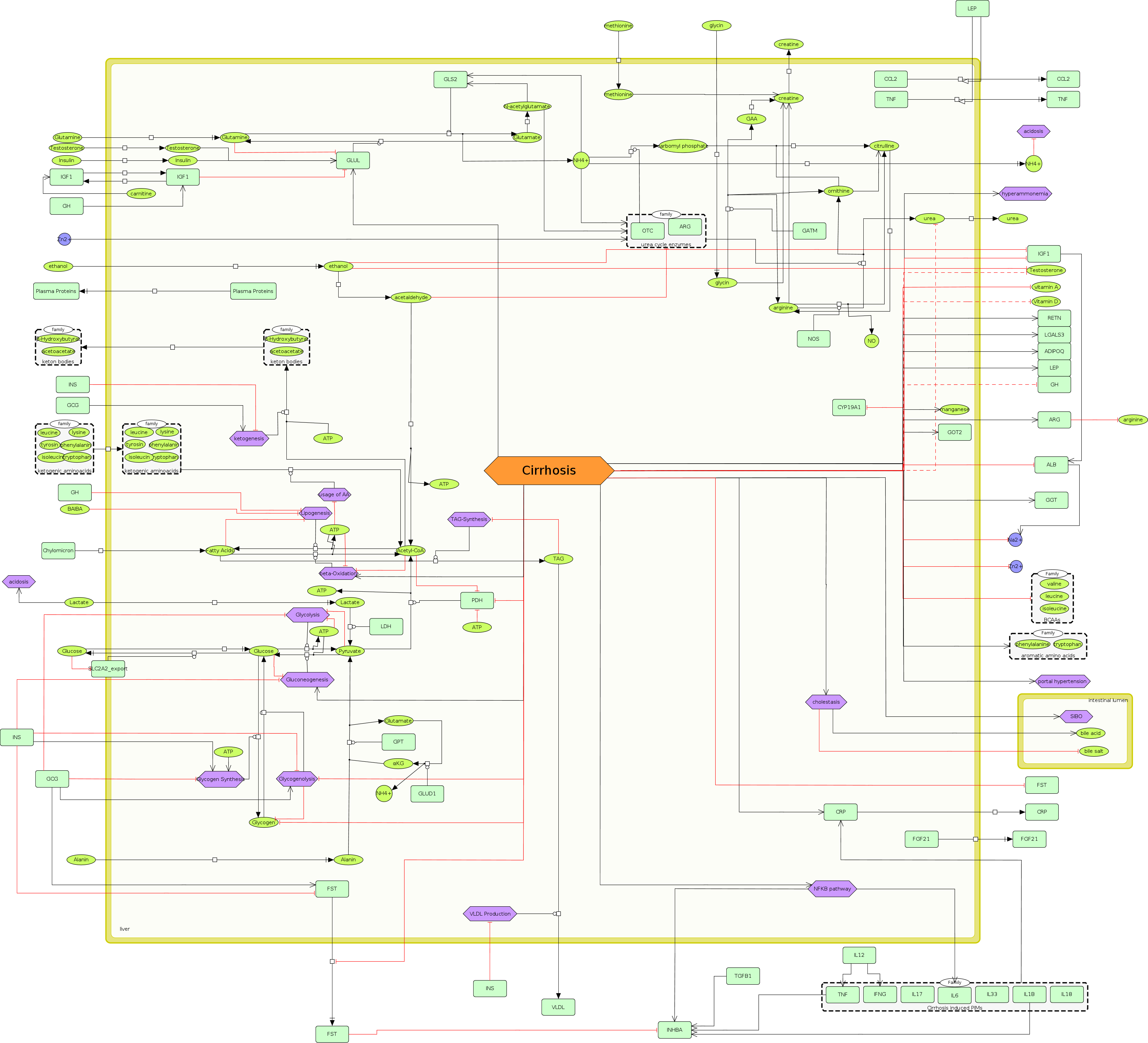

### Supplementary Figure 3

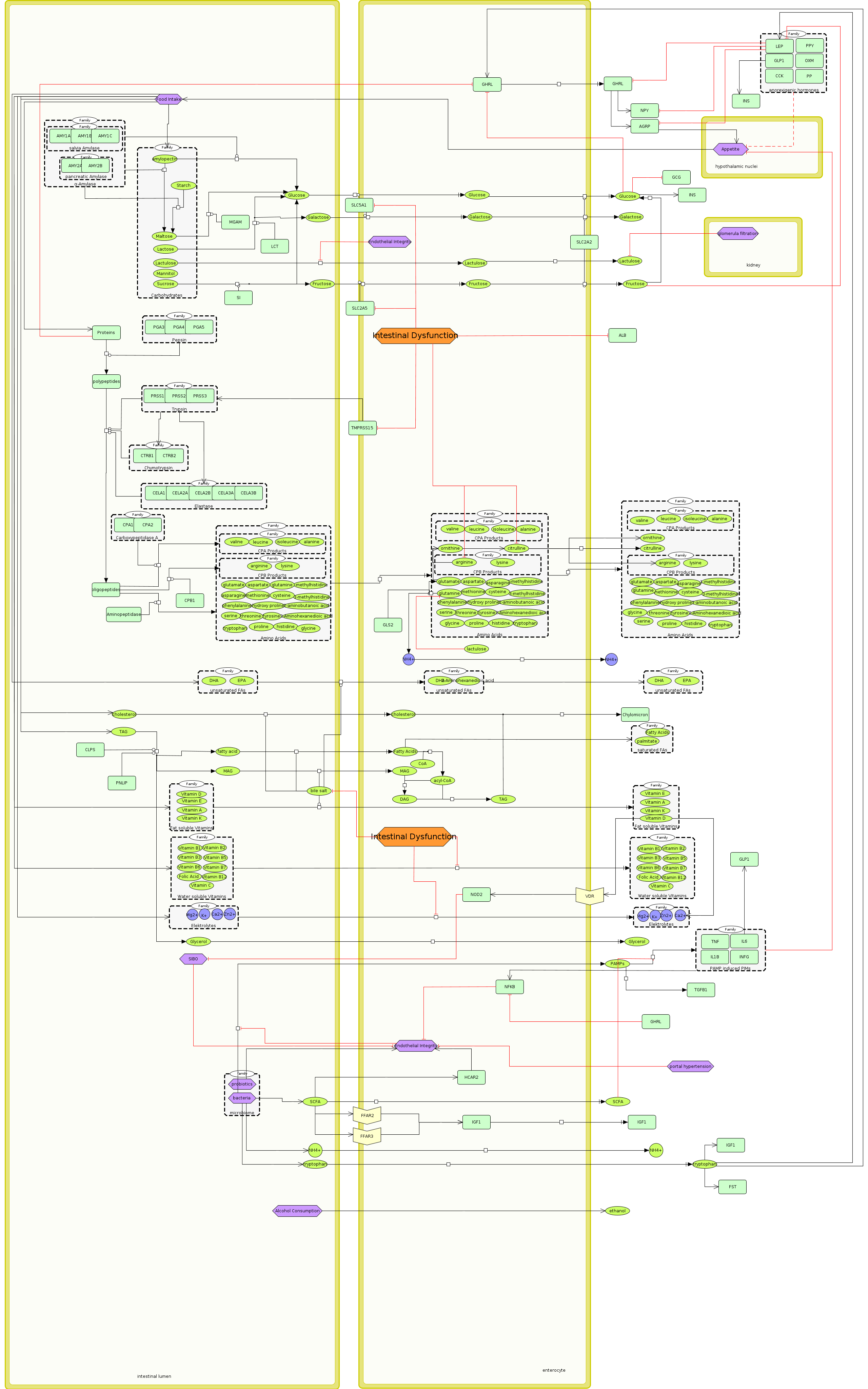

### Supplementary Figure 4

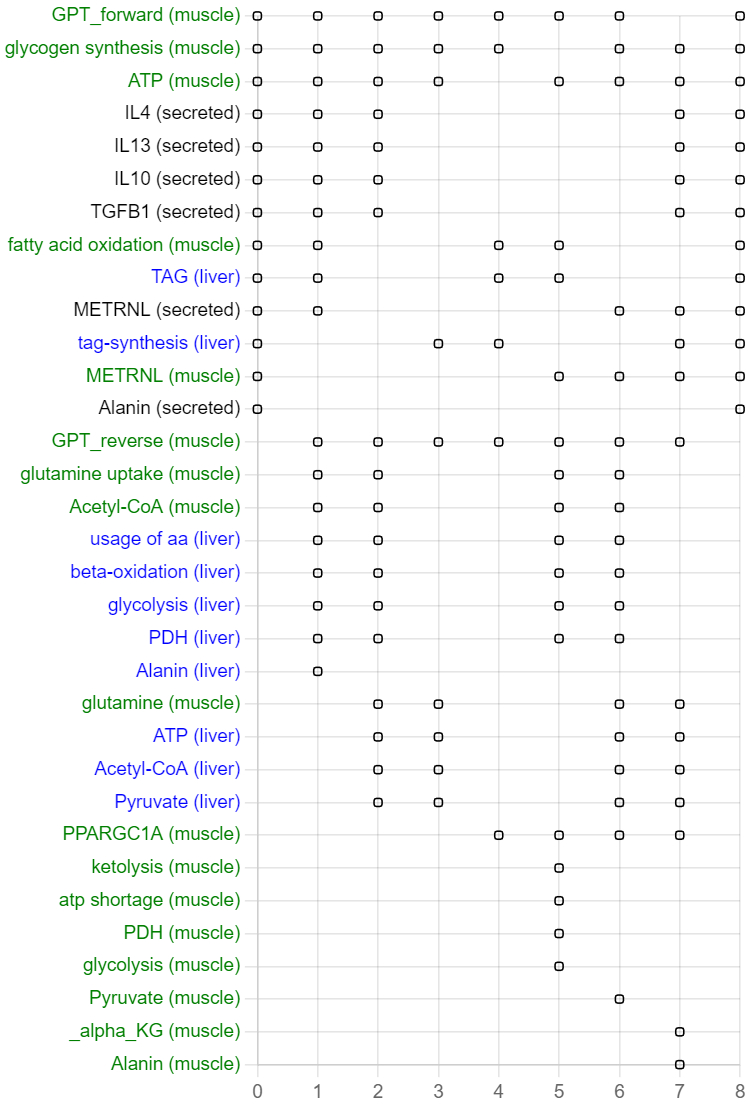
